# Acetylation of histone H2B on lysine 120 regulates BRD4 binding to intergenic enhancers

**DOI:** 10.1101/2025.02.07.637147

**Authors:** Gregory A. Hamilton, Penelope D. Ruiz, Kenny Ye, Matthew J. Gamble

## Abstract

BRD4 is a bromodomain-containing transcriptional co-regulator that plays important roles in driving transcription by binding to histone acetyl-lysines at enhancers and promoters while recruiting additional transcriptional cofactors. While the mechanisms by which BRD4 regulates transcription have been explored, the critical acetylations primarily responsible for targeting it to chromatin remain unclear. Through a machine learning approach, we determined that distinct sets of histone acetylations dominate the prediction of chromatin accessibility and BRD4 binding in distinct chromatin contexts (e.g. intergenic enhancers, gene body enhancers and promoters). Using human fibroblasts engineered to predominantly express specific histones with lysine-to-arginine mutations, we demonstrate that one such acetylation, H2BK120ac, is required to recruit BRD4 specifically to intergenic enhancers, while not affecting chromatin accessibility. Loss of H2BK120ac did not affect BRD4 binding to either promoters or gene body enhancers, demonstrating that the rules governing BRD4 recruitment to regulatory regions depends on the specific genomic context. Highlighting the importance of H2BK120ac in directing BRD4 recruitment, we found that expression of the H2BK120R mutant significantly reduces the phenotypes driven by BRD4-NUT, an oncogenic fusion protein that drives NUT midline carcinoma. This work demonstrates the critical nature that genomic context plays in BRD4 recruitment to distinct classes of regulatory elements, and suggests that intergenic and gene body enhancers represent classes of functional distinct elements.

## Introduction

Chromatin is the platform upon which gene transcription is regulated. This regulation is facilitated by the ability of chromatin to adopt distinct context-specific states at enhancers and promoters required for their function. These states are achieved through various mechanisms that alter chromatin structure including the local repositioning of nucleosomes by ATP-dependent remodeling enzymes, alterations in nucleosomal composition via the incorporation of histone variants, and the post-translational modification (PTM) of the histones. Of these mechanisms, histone PTMs are notable for the diverse mechanisms by which they contribute to transcriptional regulation. Such modifications can alter the charge of histones (as with acetylation) intrinsically changing the tightness with which they are bound to DNA, revealing, creating or masking binding surfaces for transcriptional co-regulators at promoters and enhancers^1,2^. Additionally, histone PTMs can form binding sites for recruiting specific transcriptional cofactors to chromatin. For example, bromodomain-containing factors are specifically recruited to regions of histone acetylation^3^.

Many histone PTMs are correlated with the transcriptional activity of associated genes, leading to the reasonable hypothesis that these marks play causative roles in transcriptional regulation. But often, there is little evidence to support such assertions, and in some cases the reverse causal relationship has been shown to exist. For example, maintaining the levels of H3K27ac and H3K4me3 has been shown to require ongoing transcription^4^. The difficulty in determining if a particular mark is a regulator of transcription is compounded by the fact that the writers of those marks can be promiscuous, with one enzyme catalyzing a modification at multiple positions on histones, or redundant, with multiple enzymes catalyzing the same histone PTM. The best example of both of these features comes from the histone acetyltransferases p300 and CBP which are both capable of depositing acetylatation at many lysines on histones (and non-histone proteins) and both have a high degree of overlap in their target sites^5,6^.

Enhancers play an important role in establishing cell type-specific transcriptional profiles and are sites of active chromatin remodeling and histone modification^7,8^. Enhancers have specific epigenetic profiles which distinguish them from promoters and other regulatory elements. Most notably, while both promoters and enhancers are marked by H3K27ac, enhancers are distinguished by being predominantly marked by H3K4me1 while active promoters are marked by H3K4me3^1,9^. Recently, an understudied set of acetylations on H2B were associated with active, cell type-specific enhancers^10,11^, but little is known about their causative role in the regulation of transcription.

One of the most well-studied readers of histone acetylations is BRD4, a member of the BET family of proteins which bind to acetyl-lysines through its double bromodomains^3,12^. BRD4 can bind to both enhancers and promoters^13–16^. It can promote transcription through interactions with the Mediator complex or as part of the super-elongation complex^14,17–19^. As a major regulator of transcription dysregulated in cancer, BRD4 has garnered much interest as a therapeutic target for a variety of tumor types using inhibitors that block the association of its bromodomains to acetyl-lysines^13,20–22^. These inhibitors such as JQ1, cause the rapid dissociation of BRD4 from chromatin, highlighting the importantance of acetyl-lysine binding for its correct genomic targeting. While there have been biochemical studies demonstrating which histone acetylations BRD4 can interact with, there is very little known about which acetylations are important for its targeting in cells.

Here we demonstrate that H2BK120ac acts as a critical regulator of BRD4 binding to intergenic enhancers. Mutation of H2B lysine 120 to arginine (H2BK120R), preventing its acetylation, leads to a dramatic loss of BRD4 binding specifically at intergenic enhancers, while leaving its ability to be targeted to promoters unaffected, demonstrating that the rules dictating recruitment of BRD4 depend on the larger chromatin context of the site. Highlighting the pathophysiological relevance of this finding, we show also that cell harboring the H2BK120R are protected from the effects of BRD4-NUT fusion protein expression.

## Results

### A set of H2B acetylations uniquely predict intergenic enhancer accessibility

Two key features of active enhancers are a high degree of chromatin accessibility and the deposition of specific histone marks such as H3K4me1 and H3K27ac. Histone acetylation has long been associated with enhanced chromatin accessibility at both enhancers and promoters^23^. We sought to gain a better understanding of the context-specific role of histone acetylations at distinct transcriptional regulatory regions (i.e. promoters and enhancers). To do this we made use of data from IMR90 human primary lung fibroblasts since there is both a compendium of available histone acetylation ChIP-seq data on ENCODE and published cell type-specific enhancer annotations^24–26^. We divided regulatory regions into three classes, promoters regions (N = 26,305) which we defined as a 1 kb region centered on annotated transcription start sites (TSS), intergenic enhancers (IGEs, N = 31,109) and gene body enhancers (GBEs, N = 38,033) (Fig. 1a).

**Figure 1.**
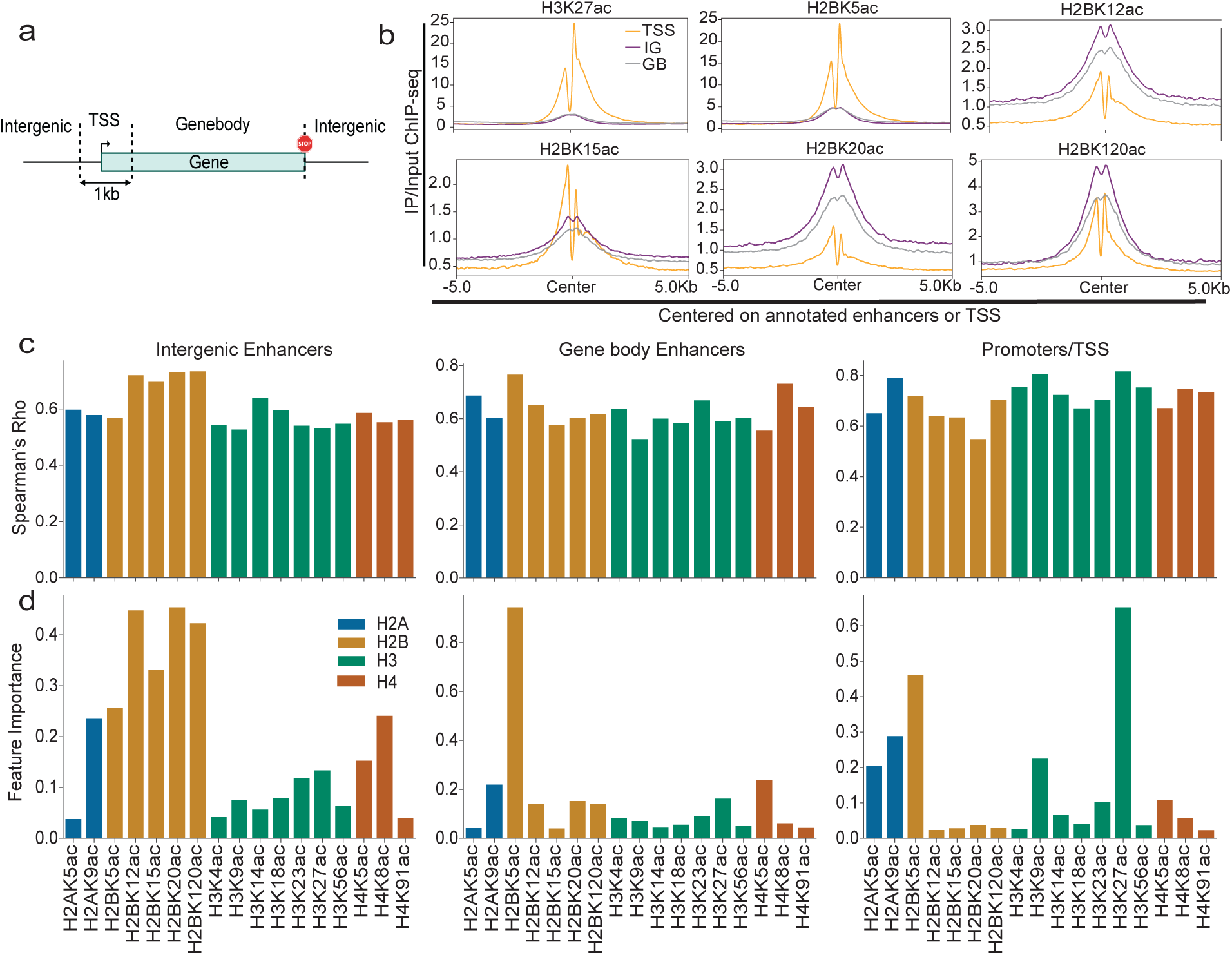
H2B acetylations uniquely mark and predict accessibility of intergenic enhancers. (a) Diagram depicting how enhancers were assigned to genomic regions. (b) Meta plots of ChIP-seq signal at promoters (orange) N=26305, gene body enhancers (grey) N = 38033 and intergenic enhancers (purple) N = 31109 (c) Spearman correlation of ENCODE IMR90 Histone acetylation ChIP-seq signal with IMR90 ATAC-seq signal at intergenic enhances (left), gene body enhancers (center) and promoters (right) (d) Random forest regressor machine learning feature importance of ENCODE IMR90 Histone Acetylation ChIP-seq signal for predicting IMR90 ATAC-seq signal at intergenic enhances (left), gene body enhancers (center) and promoters (right).

Using meta plots of the ChIP-seq data, we explored the differential enrichment of each histone acetylation across promtoers, IGEs and GBEs (Fig. 1b and S1a). This analysis revealed that while some acetylations (e.g. H2BK5ac, H3K27ac) predominantly mark promoters, other acetylations, specifically those on H2B K12, K20 and K120, predominantly mark enhancers, especially IGEs, suggesting that these H2B acetylations may play a context specific role in regulating IGE activity.

Chromatin accessibility is an important indicator of both promoter and enhancer activity. We performed ATAC-seq on IMR90 cells to determine chromatin accessibility genome wide (QC in Sup Table 1). To determine which histone acetylations are most associated with chromatin accessibility at each region of interest, we calculated the spearman correlation between each histone acetylation ChIP-seq dataset and ATAC-seq signal. While this again suggested a special role of H2B K12, K20 and K120 at IGEs, the general correlation between the various histone marks made identifying the most important marks unclear (Fig 1C).

To determine the most important acetylations for predicting chromatin accessiblity among the pool of highly correlated marks, we used a machine learning approach, random forest regression, to uncover the acetylations that conveyed the most information about chromatin accessibility at each class of regulatory region (Fig 1d). A separate model was fit for each regulatory class (e.g. promoters, GBEs, IGEs). All three models had strong R^2^ values (0.83 for promoters, 0.76 for GBEs and 0.78 for IGEs), indicating the ability to predict chromatin accessibility with high accuracy (Fig. S1b). Notably, H3K27ac, H3K9ac and H2BK5ac were the best predictors of promoter accessibility. H2BK5ac also had the highest feature importance for predicting accessibility at GBEs. Finally, acetylation of H2B at K12, K20 and K120 had a unique and dominant feature importance for predicting accessibility at IGEs. This finding is consistent with previous work documenting H2B acetylation as markers of active cell type-specific enhancers^10,11^.

### H2BK120ac contributes to IGE regulation of gene expression

The machine learning data described above led us to hypothesize that a subset of H2B acetylations contribute to enhancer function and thereby enhancer-dependent regulation of gene transcription. To test this, we engineered IMR90 cells to ectopically express tagged H2B mutants where either lysine 12 or 120 is replaced with arginine (Fig. 2a). As a control, we generated IMR90 cells ectopically expressing tagged wild-type H2B. Importantly, the ectopically expressed H2B were expressed at a high enough level to out compete endogenous H2B, to become the predominant H2B expressed in these cells. We have used this method previously to explore the function of other H2B acetylations^27^.

**Figure 2.**
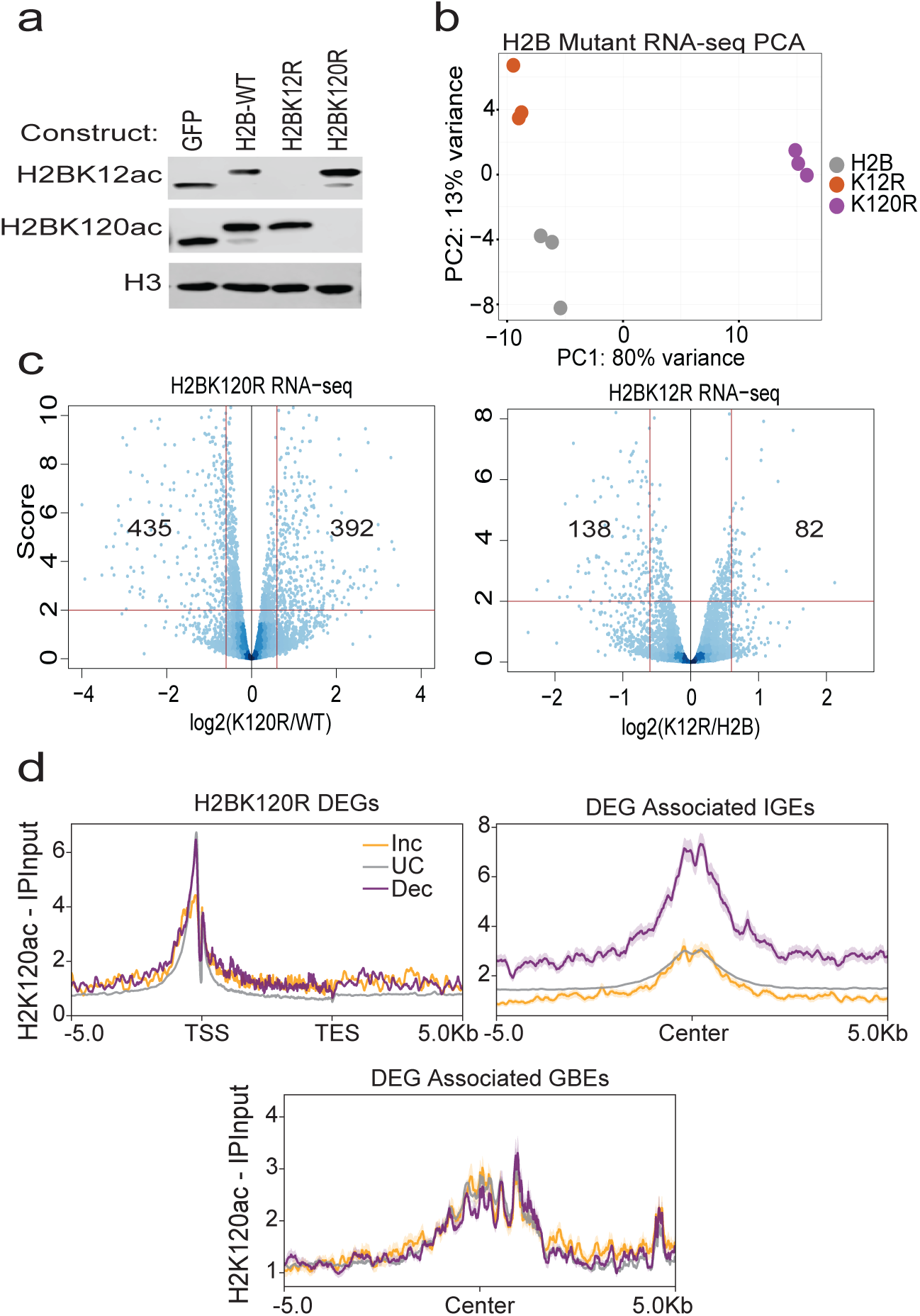
H2BK120ac regulates gene expression through intergenic enhancers. (a) Western blot of depicting H2BK12ac and H2BK120ac in the ectopic H2B mutant lines. (b) PCA plot of RNA-seq data from the ectopic H2B Mutant lines (c) Volcano plots of H2BK12R (left) and H2BK120R (right) RNA-seq data. Y-axis depicts score (log_10_(pValue)) and x-axis depicts Log_2_(Exp/Control). (d) Meta plots of H2BK120ac across DEG gene bodies (left), associated IGEs (right, N = 1084(Increased),17009(UC),1503(Decreased)) and GBEs (bottom, N = 814(Increased),14256(UC),1234(Decreased)).

RNA-seq was used to determine the effect of each mutant on gene transcription (Quality controls are documented in Sup Table 1). Principal component analysis (PCA) demonstrated that the replicates segregated by cell line, with PC1 (80% of variance) representing changes due to H2BK120R expression and PC2 (13% of variance) representing changes caused by expression of H2BK12R (Fig. 2b). Using a log_2_ fold change cutoff of 0.6 and an adjusted p value of 0.01, we identified only 212 differentially expressed genes (DEGs) upon replacement of H2B with H2BK12R (138 decreased, 82 increased) (Fig. 2c). More extensive changes in gene expression were seen upon replacement of H2B with H2BK120R (827 DEGs, 435 decreased and 392 increased) (Fig. 2d). We performed GSEA analysis on the DEGs for both datasets and found an enrichment for cell identity related pathways for the H2BK120R DEGs (Fig. S2a,b).

To determine if promoter levels of H2BK120ac are predictive of regulation upon H2BK120R expression, we determined the average level of H2BK120ac at promoters of genes with increased, decreased or unchanged expression upon H2BK120R expression, and found no significant difference (Fig 2d). This suggested that the promoter was not necessarily the critical site of action for H2BK120ac to regulate gene expression, so we turned our attention to enhancers. We identified the closest enhancer to each genes promoters and then examined the average level of H2BK120ac at enhancers associated with genes with increased, decreased or unchanged in cell expressing H2BK120R. We found that genes with decreased expression in the H2BK120R cells have significantly higher levels of H2BK120ac at their nearest IGE (Fig 2e,right) while H2BK120ac levels at promoters and the nearest GBE were indistinguishable from genes that did not change in expression. Overall, this data demonstrates that H2BK120ac supports the function of IGEs to promote transcription of their associated genes.

### H2B acetylation on K12 and/or K120 does not generally promote chromatin accessibility at IGEs

Active enhancers maintain a high degree of chromatin accessibility allowing transcription factors and other co-regulators to gain access to their binding sites. Given that H2BK120ac was one of the best predictors of IGE chromatin accessibility, we hypothesized that H2BK120ac functionally promotes an open chromatin structure at enhancers. To determine if H2BK120ac contributes to chromatin accessibility, we performed ATAC-seq on IMR90 cells expressing ectopic H2BK12R, H2BK120R or wild-type H2B as a control. In the control cells, we identified 185,212 peaks of accessibility with sufficient coverage to determine statistical differences between conditions. PCA analysis demonstrated good separation between H2BK120R samples and H2B wild-type expressing cells. However, H2BK12R cells did not separate from H2B wild-type in any principal component explored, suggesting that H2BK12ac contributes minimally to changes in accessibility. Differentially accessible regions (DARs) were determined using DESeq2 with cutoffs of 0.001 for p-value and an absolute log_2_ fold change greater than 1. Using these cutoffs we determined that the H2BK12R cells had a minimal effect on accessiblity with less than 30 total peaks having significant differential accessibility (Fig. S3a). The H2BK120R cells, on the other hand, led to widespread changes in accessiblity with 5552 peaks decreasing and 3081 peaks increasing in accessibility compared to the wild-type H2B-expressing control cells (Fig. 3b).

**Figure 3.**
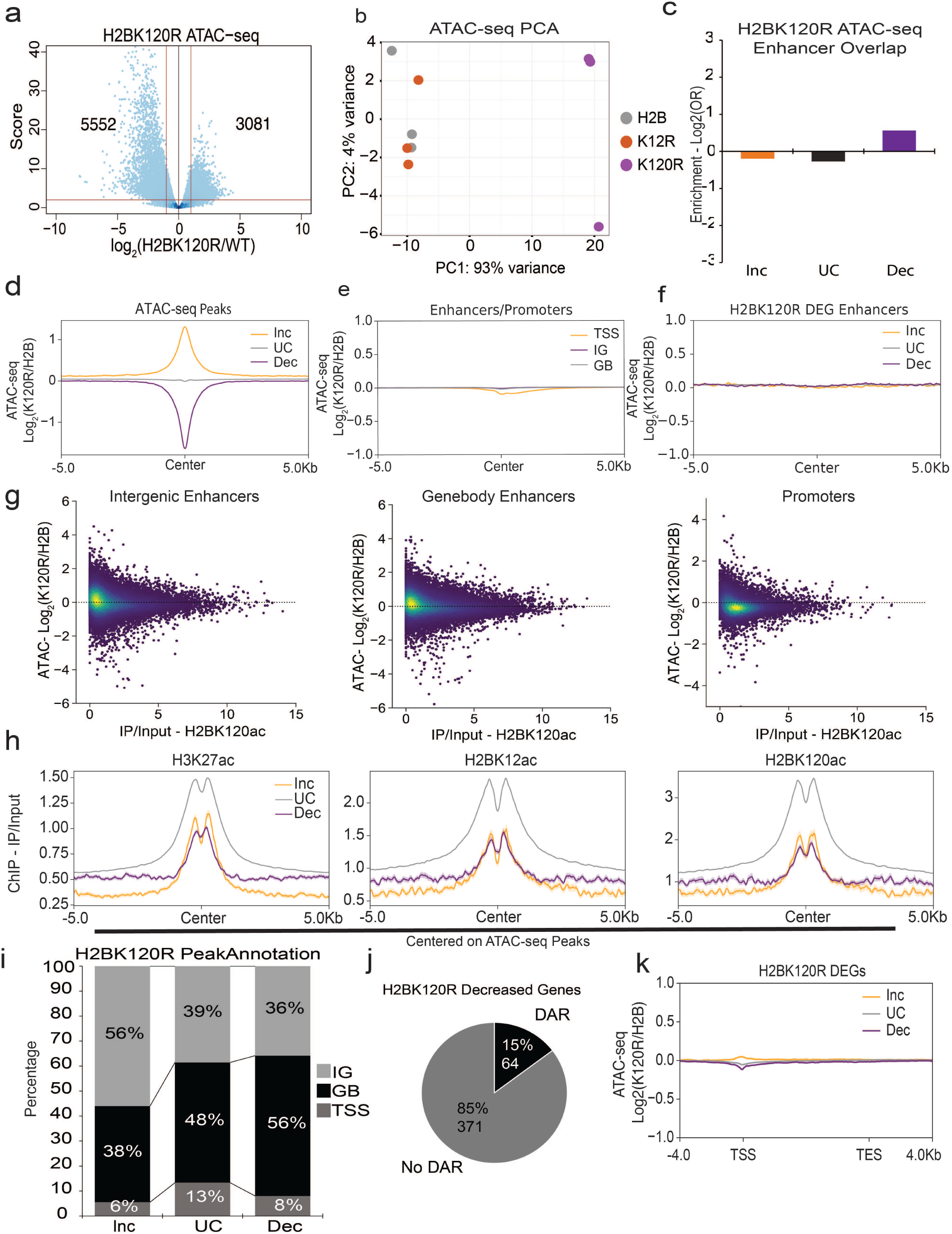
H2BK120ac does not regulate accessibility of intergenic enhancers. (a) Volcano plot of H2BK120R/H2B WT ATAC-seq data. Y-axis depicts score (log_10_(pValue)) and x-axis depicts Log_2_(Exp/Control). (b) PCA plot of ATAC-seq data from the ectopic H2B Mutant lines. (c) Bar plot depicting Log_2_(OR) of a fisher exact test for overlap between ATAC-seq DARs and enhancers. (d) Meta plot of Log_2_(H2BK120R/H2B WT) ATAC-seq signal at the H2BK120R ATAC-seq DARs. Increased (N=3081), Unchanged (N=176579) and Decreased (N=5552). (e) Meta plot of Log_2_(H2BK120R/H2B WT) ATAC-seq signal TSS (orange), gene body enhancers (grey) and intergenic enhancers (purple). Same regions used in Figure 1B. (f) Meta plot of Log_2_(H2BK120R/H2B WT) ATAC-seq signal at H2BK120R DEG associated enhancers. Increased DEG (orange), Unchanced DEG (grey) and Decreased DEG (purple). (g) Scatter plots of H2BK120ac ChIP-seq (x-axis) and Log_2_(H2BK120R/H2B WT) ATAC-seq (y-axis) at intergenic enhancers (left), gene body enhancers (middle) and promoters (right). (h) Meta plots of H3K27ac (left), H2BK12ac (middle) and H2BK120ac (right) ChIP-seq signal at H2BK120R ATAC-seq DARs. (i) H2BK120R ATAC-seq DAR annotation using the rules from figure 1A. (j) Pie chart of H2BK120R decreased DARs with H2BK120R decreased DEGs. (k) Meta plots of log_2_(H2BK120R/H2B) ATAC-seq at H2BK120R DEGs.

We next looked for an enrichment between IGEs and DARs induced by H2BK120R expression, finding only a modest but not statistically significant enrichment for deacreased DARs in IGEs (Fig. 3c). While we expected a strong association between IGEs and DARs caused by H2BK120R expression, we were surprised to find only a modest enrichment that did not pass a p-Value<0.05 cutoff for statistical significance (Fig. 3c). Meta plots of ATAC-seq signal at promoters, IGEs and GBEs, when compared to the signal seen at DARs, further demonstrate the lack of significant changes in accessibility at these regulatory regions (Fig. 3d,e). Next, we determined the average change in accessibility at the IGEs associated with H2BK120R DEGs and again found no meaningful difference (Fig. 3f). Additionally, we found no relationship between the change in chromatin accessibility in the H2BK120R cells and the level of H2BK120ac at neither IGEs, GBEs nor promoters (Fig. 3g). Finally, H2BK120ac, along with other enhancer marks were actually depleted from increased and decreased H2BK120R associated DARs (Fig. 3h). Meta plots for the rest of the histone acetylation ChIP-seq datasets at DARs indicate that no histone acetylations were enriched for decreased or increased DARs (Fig. S3b) From all of this, we conclude that H2BK120ac does not promote IGE function by mediating the accessibility of enhancers.

Given the lack of connection between DARs in H2BK120R cells and sites of H2BK120ac, we were left with the question of what does dictate where chromatin accessibility is altered in the H2BK120R cells. H2BK120 is not only a site of acetylation but also mono-ubiquitylation^28,29^. Importantly, H2BK120ub1 has a localization pattern that is highly distict from H2BK120ac; whereas H2BK120ac predominantly marks intergenic enhancers, H2BK120ub1 is deposited co-transcriptionally within active genes^28,29^. We hypothesized that the changes in chromatin accessibility seen in the H2BK120R cells are due to loss of H2BK120ub1. Consistent with this, DARs with decreased accessibility in H2BK120R cells are enriched for gene bodies (Fig. 3i). This suggests that the changes in chromatin accessibility seen in gene bodies in H2BK120R cells is due to loss of H2BK120ub1 and not H2BK120ac. Finally, only 15% of H2BK120R DEGs had a H2BK120R DAR and H2BK120R DEGs only had minor accessibility changes localized to their promoter regions (Fig 3j,k). Together, these data indicate that the gene expression changes seen in H2BK120R are not due to changes in chromatin accessiblity. Yet, the level of H2BK120ac at decreased gene associated IGE suggests that H2BK120ac regulates gene expression through IGE activity just not IGE accessibility.

### Acetylation of H2B is associated with BRD4 recruitment to IGEs

Given that loss H2BK120ac did not lead to changes in chromatin accessibility at IGEs, we decided to turn our attention to the possibility that this mark may regulate IGE function by promoting the recruitment of an enhancer binding cofactor. Specifically, we turned our attention to BRD4 for two reasons. First, we found a strong overlap between genes downregulated in H2BK120R cells and genes downregulated upon treatment of IMR90 cells with JQ1^30^ (Fig. 4a). In addition, data from a previously published peptide array demonstrated that H2BK120ac had high affinity for the second bromodomain of BRD4, along with a variety of other acetylated positions^3^. The genes downregulated by both JQ1+ and H2BK120R (N = 225) were analyzed via GSEA and found to be enriched for cell identity and signaling pathways, notably epithelial to mesenchymal transition pathway. This again points to their shared function being through enhancers as genes tied to cell identity are typically regulated by enhancers^8^.

**Figure 4.**
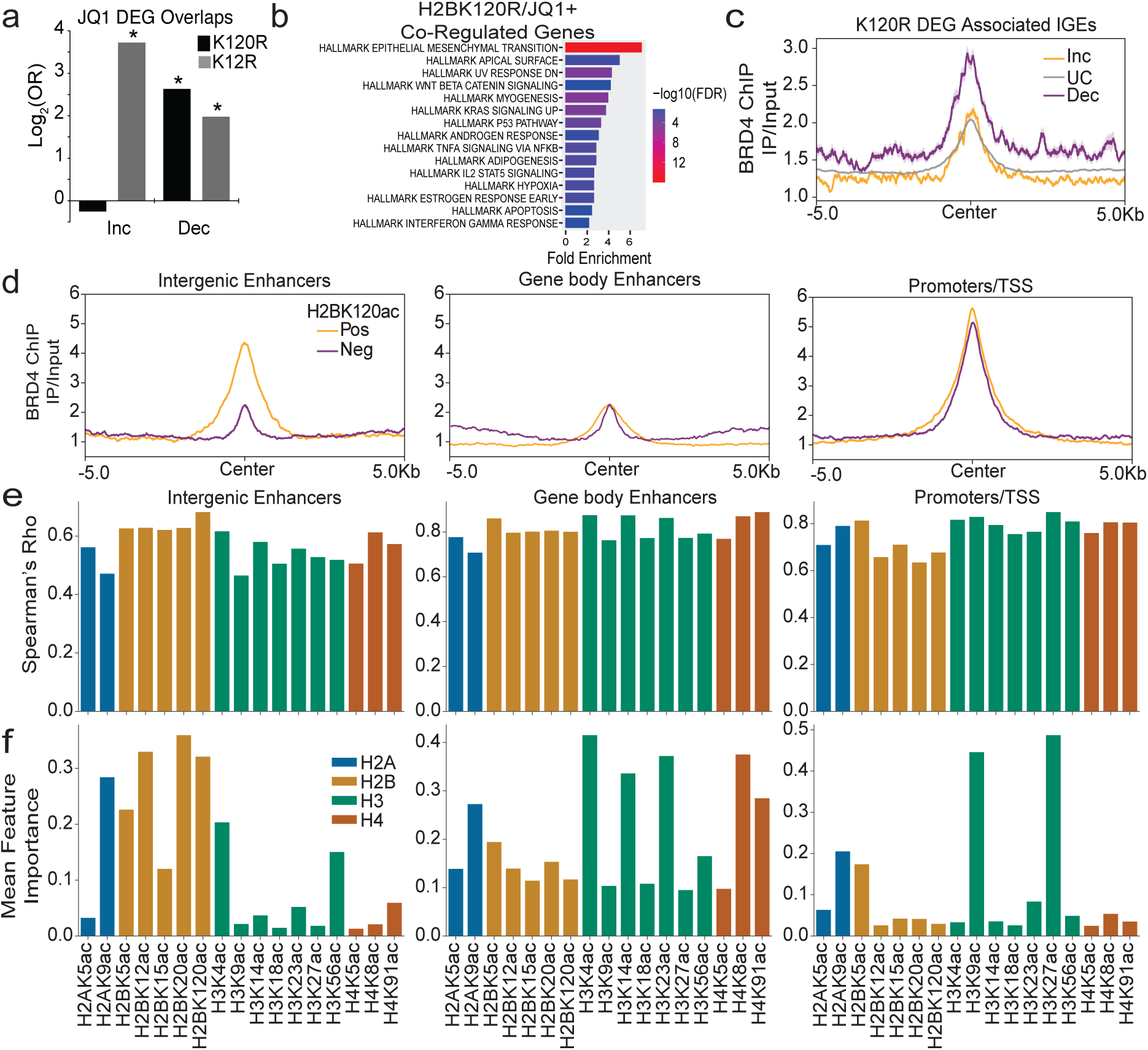
H2B acetyls predict BRD4 binding at intergenic enhancers and BRD4 binding regulates a significant number of the same genes as H2BK120R. (a) Bar plot of Log_2_(Odds ratio) from a fisher exact test overlapping JQ1 and H2BK120R DEGs. (*p<0.00001, Fisher’s Exact Test) (b) GSEA Analysis of shared JQ1 and H2BK120R DEGs. N = 225(c) Meta plot of BRD4 ChIP-seq signal at H2BK120R DEG associated enhancers. (d) Meta plot of BRD4 ChIP-seq signal at H2BK120ac positive(orange) and negative (purple) IGEs (left,10224 positive, 14547 negative), GBEs (middle, 10224 positive, 17547 negative) and promoters (right, 4322 positive, 4930 negative). (e) Spearman correlation of ENCODE IMR90 histone acetylation ChIP-seq signal with BRD4 ChIP-seq signal at IGEs (left), GBEs (e) and promoters (right). (f) Random forest regressor machine learning feature importance of ENCODE IMR90 histone acetylation ChIP-seq signals predicting BRD4 ChIP-seq signal at IGEs (left), GBEs (e) and promoters (right).

To explore the relationship between BRD4 and H2BK120ac, we performed ChIP-seq for BRD4 in our control cells and found that it is enriched at enhancers associated with genes downregulated in H2BK120R cells (Fig. 4c). Next, we subset IGEs, GBEs and promoters into those that are positive for H2BK120ac (Log_2_(IP/input) > 0.5) or negative for the mark. We found that BRD4 was specifically enriched at IGEs when they were marked by H2BK120ac (Fig. 4d). Interestingly, H2BK120ac status does not seem to differentiate BRD4 binding at either GBEs or promoters.

To get a more agnostic view into the histone acetylations that serve as binding determinants for BRD4, we determined the correlation of each histone PTM to BRD4 at IGEs, GBEs and promoters. Similar to what we found for chromatin accessibility in Fig. 1, we saw that most histone acetylations correlated with BRD4 binding regardless of element type (Fig 4e). So again we turned the machine learning approach of random forest regression to determine which histone acetylations were the predominant predictors of BRD4 biniding. We found that distinct groups of acetylations have high feature importance at each type of regualtory region (Fig. 4f). H3K9ac and H3K27ac were the best predictors of BRD4 binding to promoters, while acetylation on H3 at lysines 4, 14, and 24 and H4K8ac were strong predictors of BRD4 binding to GBEs. Finally, we found that H2B acetylations, specifically at lysines 12, 20 and 120, had the highest feature importance for BRD4 binding to IGEs. Quality controls for all three models can be found in Fig. S4. Together, this suggests that different histone acetylations may contribute to BRD4 binding in a manner that depends on the type of regulatory element (e.g. IGEs, GBEs or promoters) and that H2BK120ac may play an important role at IGEs.

### H2BK120ac regulates BRD4 binding to IGEs, but not GBEs or promoters

To test the hypothesis that H2BK120ac may be required to recruit BRD4 to a large subset of IGEs, we performed ChIP-seq in our H2BK120R expressing cells. Importantly, expression of H2BK120R and the concomitant loss of H2BK120 acetylation did not result in downregulation of BRD4 protein (Fig. 5a). Strikingly, we found that H2BK120R expression led to a loss of BRD4 binding specifically at IGEs, while average levels BRD4 at promoters and GBEs remain unchanged (Fig. 5b). H2BK12R expression had no effect on BRD4 at any of the three classes of regulatory regions explored (Fig. S5b). We found a strong association between loss of BRD4 in H2BK120R cells and the level of H2BK120ac at IGEs (Fig 5c, top-right). This association was not seen at promoters or GBEs (Fig. 5c, bottom), demonstrating that H2BK120ac is a specific determinant of BRD4 binding at intergenic enhancers. Indeed, even if we look across all BRD4 peaks, we can see that H2BK120ac is a major predictor of BRD4 loss in H2BK120R cells, highlighting the important role of H2BK120ac in mediating BRD4 binding to a subset of it’s genomic binding sites (Fig. 5c, top-left).

**Figure 5.**
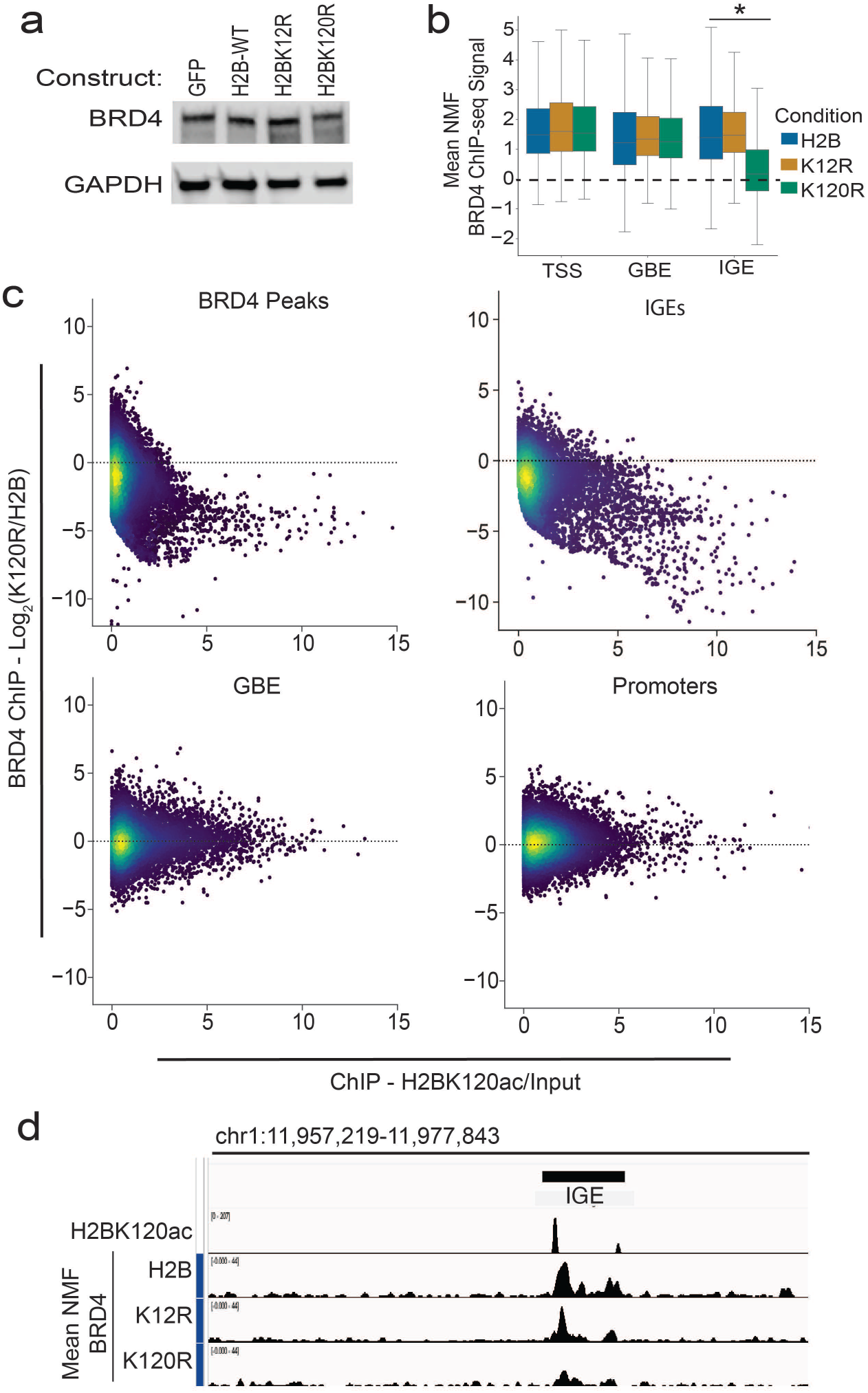
H2BK120ac regulates BRD4 binding at intergenic enhancers. (a) Western blot for BRD4 in the H2B Mutants. (N = 1) (b) Boxplot of replicate mean NMF normalized BRD4 ChIP-seq signal at promoters (N = 23845), GBEs(N = 34474), and IGEs(N = 29707). (*,pValue < 0.01,two-tailed Student’s T-test) (c) Scatter plots of Log2(K120R/H2B)(Left) NMF normalized BRD4 ChIP-seq signal (y-axis) at BRD4 peaks(N = 56725, top left), IGEs(N = 8484, top right), GBEs(N = 9729, bottom left) and promoters(N = 15281, bottom right) with IP/Input H2BK120ac ChIP-seq data x-axis. (d) IGV Plot depicting an IGE where BRD4 binding is lost in the H2BK120R mutant.

### Loss of H2BK120ac renders cells resistant to the BRD4-NUT oncogenic fusion protein

BRD4-NUT is an oncogenic fusion protein created through a translocation event in NUT midline carcinomas between the NUTM1 gene and BRD4^21,31,32^. The BRD4 portion of the fusion retains the ability to bind acetyl-lysines while the NUTM1 portion enables recruitment of p300 and CBP, acetyltransferases which can mediate the acetylation of many positions on histone, including H2B at K120^5,6,33^. With the combination of these two abilities, BRD4-NUT expression causes the formation of large acetylated “megadomains” that can span large portions of chromosomes^34^. These megadomains cause large scale chromatin disorganization^32,34^. While BRD4-NUT promotes the growth of NUT midline carcinomas, the havoc they wreak on chromatin is toxic to most cell lines^32^. Given the importance of H2BK120ac in mediating BRD4 binding to IGEs, we set out to test if H2BK120R expression can ameliorate the aberrant function of BRD4-NUT.

We generated tagged wild-type H2B, H2BK12R and H2BK120R IMR90 cells harboring a doxycycline-inducible HA-tagged BRD4-NUT transgene incorporated through lentiviral transduction. After 24 hours of induction with doxycycline we observed robust expression of BRD4-NUT (Fig. 6a). BRD4-NUT expression typically leads to an increase in the level of H3K27ac^34^, which is what we observe upon induction of BRD4-NUT (Fig. 6a,b). In addition we also observed that BRD4-NUT leads to an increase in H2BK120ac. Interestingly, the BRD4-NUT-induced increase in H3K27ac levels were abrogated in the H2BK120R expressing cells (Fig. 6a,b). Together, this suggests that H2BK120ac is upstream of H3K27ac in the BRD4-NUT-induced feed-forward loop that drives megadomain formation.

**Figure 6.**
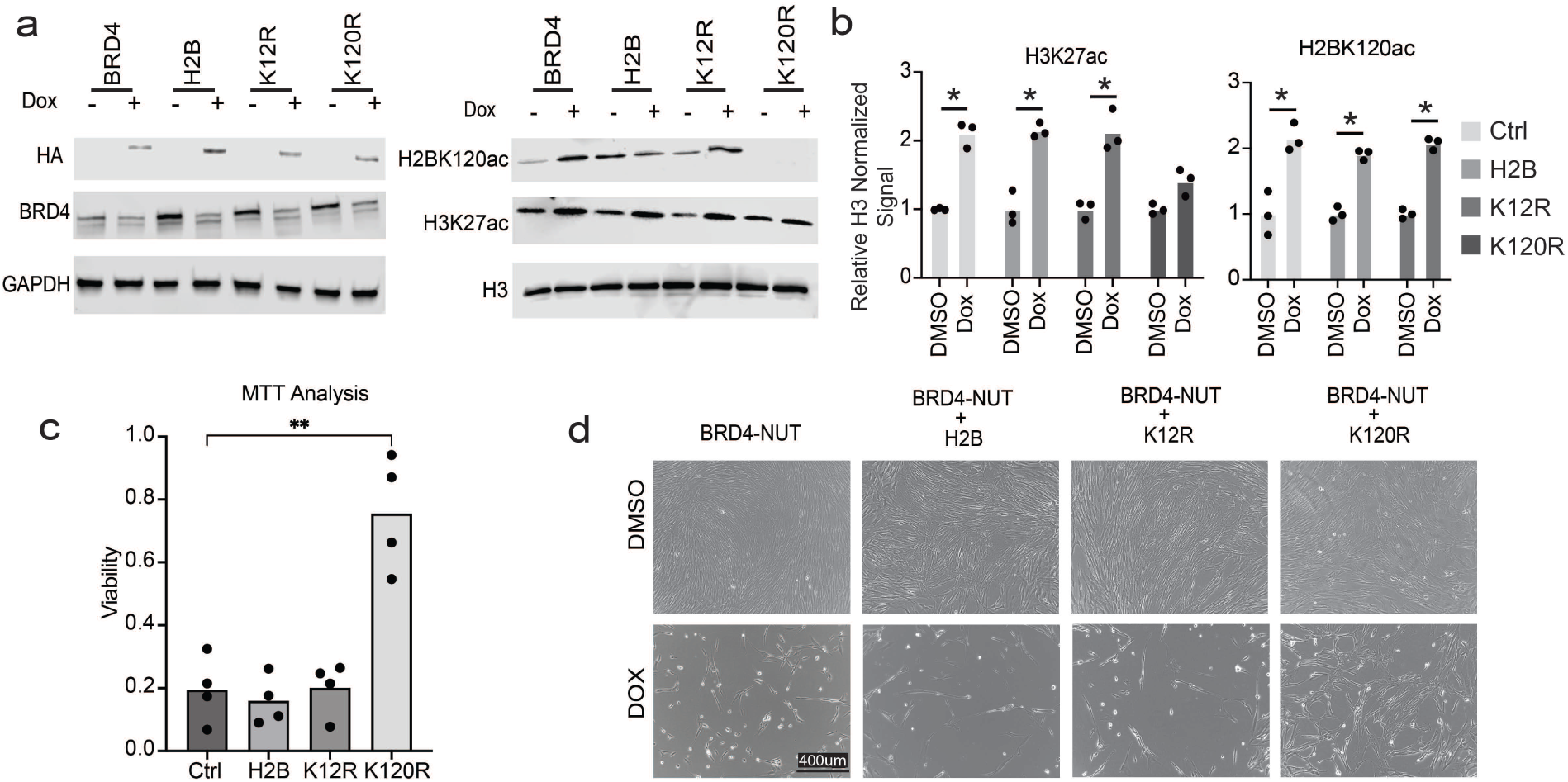
H2BK120R protects against BRD4-NUT expression driven pheontypes. (a) H2B mutant/BRD4-NUT cell line western blots for HA, BRD4, H2BK120ac, H3K27ac, H3 and GAPDH twenty-four hours post doxycycline or DMSO treatment. (b) Relative H3 normalized western blot signal for H3K27ac (left) and H2BK120ac (right) in BRD4-NUT H2B mutants in DMSO (black) and doxycycline treated (grey). (N = 3) (*,p < 0.005,two-tailed Student’s T-test) (c) MTT assay viability plot for forty-eight hours post doxycycline treatment in the inducible BRD4-NUT H2B Mutants. (**,p < 0.01, two-tailed Student’s T-test). (d) Bright field images of BRD4-NUT H2B mutants forty-eight hours after DMSO (Control/Top Row) or doxycycline (Bottom Row) treatment.

Unlike NUT midline carcinoma cells, most cell lines cannot tolerate BRD4-NUT expression^32^. Strikingly, we find that expression of H2BK120R in IMR90 cells protects them from the toxicity associated with BRD4-NUT expression as measured by an MTT assay (Fig. 6c,d). Overall, our results show that H2BK120ac plays a critical role in the epigenetic changes associated with BRD4-NUT expression, suggesting that IGEs fuction as critical nucleation sites for the formation of megadomains.

## Discussion

Genomic context plays a pivotal role in establishing the chromatin environment required for the regulation of distinct regions such as promoters, silencers, insulators, and enhancers. Our findings suggest that gene body and intergenic enhancers represent distinct functional subclasses. Our work demonstrates that H2BK120ac is critical for the recruitment of BRD4 to intergenic enhancers. This is a distinguishing feature of intergenic enhancers that differentiates them from both gene body enhancers and promoters where BRD4 recruitment is occurs independently of this PTM. Furthermore, we demonstrate the functional consequence of loss of H2BK120ac-mediated BRD4 recruitment to intergenic enhancers on transcriptional regulation.

Much of what is known about the function of particular histone PTMs at genomic regions has been inferred from associative bioinformatic approaches such as correlation and linear regression. Yet these results leave the directionality and causality of the relationship these histone marks with transcription unresolved. Through these associative approaches it had been long thought that H3K27ac^35^ is important for driving transcription at promoters but two recent studies have called it’s role into question. The first found that transcription itself is required for maintaining H3K27ac^4^ and that second found that loss of H3K27ac did not broadly effect transcription^36^. These results highlight the importance perturbation experiments for determining the functional consequences a histone PTM has (if any) on transcriptional regulation. Sankar et al. employed the “gold standard” for investigating the causitive role of particular histone modification by mutating all 28 mouse H3 alleles to replace lysine 27 with an arginine^36^. While clearly heroic, this difficult and time consuming approach is currently beyond the capabilities of most labs in the chromatin field. We used a simpler approach here and in the past, ectopically expressing tagged histone mutants at a level that replaces the overwhelming majority of the endogenous histone in chromatin^27^. Building on previous work which found H2BK120ac/H2B acetylations are highly associated with and predictive of cell type specific enhancers^10,11,35^, this approach allowed us to determine that H2BK120ac regulates BRD4 binding to intergenic enhancers.

Work investigating BRD4 binding to chromatin has been heavily reliant on the use of bromodomain inhibitors and biochemical peptide-based assays^3,12,20,30^. Bromodomain inhibitors prevent BRD4 binding globally, and therefore, cannot provide information about the role of context-specific BRD4 binding. Biochemical peptide based assays use the bromodomains of BRD4 in lieu of the full length protein and evaluate binding in the context of short acetylated peptides of histone tails^3,37^. Through these approaches the field has been able to uncover the plethora of histone acetylations BRD4’s bromodomains are capable of binding and the mechanisms through which it regulates gene expression, the recruitment of mediator at enhancers and the recruitment of the super elongation complex at both enhancers and promoters^12,14^. Our work demonstrates the first evidence of a particular histone PTM (H2BK120ac) being required for BRD4 binding to a subset of chromatin in living cells and its subsequent affect on gene transcription.

BRD4 regulates transcription through at least two distince mechanisms, the recruitment of mediator at enhancers and the recruitment of the super elongation complex at both enhancers and promoters^12,14^. Recent work has shown that not all of BRD4’s roles in transcriptional regulation require its bromodomains. BRD4-mediated promoter-proximal pause release is independent of its ability to bind acetylated histones^38^. Given the dependence of BRD4 on H2BK120ac for its binding to intergenic enhancers, we speculate that BRD4 recruited to these sites is therefore less likely to function in promoter-proximal pause release and instead is more consistent with the recruitment of the mediator complex.

We show that the binding of BRD4 to gene body enhancers is independent of H2BK120ac. But, why? One potential explaination is that H2B K120 is typically ubiquitylated in the transcribed regions of active genes. H2BK120ub1 plays important roles in promoting H3K79me2, a mark involved in regulating transcriptional elongtation^28,29,39^. This leaves open the possibility that other histone acetylations are responsible for BRD4 recruitment to this enhancer subclass. Our machine learning results indicate that H2BK9ac, H3K4/9/14/23/27ac, and H4K8/91ac are all candidates worth pursuing. In the future, our ectopic histone mutant driven approach may be able to determine which marks regulate BRD4 binding to gene body enhancers. Additionally, all of these marks can be bound by one or both of BRD4’s bromodomains^3,37^.

BRD4 has become a major target for cancer therapeutics^13,40^ due to its role in driving oncogensis in a variety of cancers^30,41^. A particularly deadly BRD4 mutation drives NUT midline carcinomas^32^. Through a translocation event, BRD4 and NUTM1 form a fusion protein (BRD4-NUT) that creates large acetylated chromatin domains leading to wide scale genomic disorganization^32,40^. We found that loss of H2BK120ac blunted the BRD4-NUT-dependent increase in H3K27ac and additionally blocked BRD4-NUT-mediated cytotoxicity. Loss of BRD4-NUT’s ability to bind at intergenic enhancers may reduce its overall ability to nucleate hyperacetylated megadomains and/or slow their ability to spread. Futher work is required to determine if such a mechanism is important in NUT midline carcinoma cells.

Our results demonstrate that the rules which dictate BRD4 binding to chromatin depend on the genomic context. We show that H2BK120ac is critial for the binding of BRD4 to a subset of enhancers, specifically those found outside of gene bodies. While at the same time we demonstrate that this acetylation is despensible for the binding of BRD4 to gene body enhancers or promoters.

## Methods

### Machine Learning model

We created a Random Forest Regressor model utilizing the python Scikit-Learn module. The model utilizes ENCODE ChIP-seq signal data(Supp Table 2) as features and either ATAC-seq signal or BRD4 ChIP-seq signal as the target.

The ENCODE ChIP-seq data served as the features or independent variables while the ATAC-seq or BRD4 ChIP-seq signal served as the dependent variable. The Random Forest Regressor model was set up with 10,000 initial estimators, a minimum sample split size of ten and five for the minimum samples per leaf. We set aside five percent of the total data to be used for post model generation testing and used the remaining ninety-five percent for training the model. The testing data was then used to predict target signal and we compared predict vs actual to determine the R-squared and mean squared error of the model.

We utilized the Scikit-learn permutation importance function to run 100 permutations of the model to ensure that we results were stable and repeatable. The model code is available on our lab github page: https://github.com/TheRealGambleLab.

### Cell Lines

All the reported work was done using IMR90 human lung fibroblasts that have been hTERT immortalized. Cells were cultured following ATCC guidelines using MEM with 10% FBS. H2B mutant lines were established using lentivirial mediated ectopic expression of wild type H2B, H2B with lysine 12 or lysine 120 substituted with an arginine. Two infections were required to fully replace the endogenous H2B and the lines were selected for using Puromycin and Neomycin. Selection was done for 48 hours prior to experiments and puromycin/neomycin was removed when plating for experiments. The puromycin resistant construct H2Bs are tagged with FLAG and the neomycin resistant construct H2Bs are tagged with HA. The constructs were generated using the pLVX-IRES vector with cloning methods following our previously published protocol^27^. Doxycycline inducible BRD4-NUT lines were created using a previously published construct provided by the French lab^34^. Transfection and infection for all lines was done following our previously published protocol^27^. Cells were selected with puromycin(.9ug/mL), neomycin (.5mg/mL) and/or blasticidin (3ug/mL) for 48 hours post infection and kept in selection media for passaging plates while non-selection media was used when plating for experiments. All cells were collected at 80-90% confluency for all experiments. BRD4-NUT induction was performed using 1 ng/mL of doxycycline 24 hours prior to collection for the western blots and 48 hrs prior to analysis for the MTT assay^42^.

### Immunoblotting

10 cm plates were harvested using cold PBS and a cell scraper. The resultant was then spun down and the supernatant was removed. The cell pellet was lysed with a buffer containing 25mM Tris, 150mM NaCl, 0.1 mM EDTA, 10% Glycerol, and 0.1 mM Triton X-100. Protease inhibitor and 1 mM DTT were added to the buffer prior to use. The lysate was then cleared by centrifugation at 14,000 rpm for ten min. Using the supernatant, whole cell extract, protein concentrations were calculated by a Bradford assay (company and cat number). The remaining pellet containing the chromatin was then acid extracted by resuspending it in 1 M HCl for two hours. The mixture was then spun down at 14000 rpms for five minutes and the supernatant was mixed with 2M Tris resulting in a final mixture of 80% 1M HCl and 20% 2M Tris. Whole cell extracts and acid extracts were subjected to SDS-PAGE followed by immunoblotting. The whole cell extracts were immunoblotted overnight at 4C with antibodies: BRD4 (Cell Signaling; E2A7X), HA (Cell Signaling; C29F4) and GAPDH (Cell Signaling; 14C10). While the acid extracts were immunoblotted with H2BK12ac (Active Motif; 39669), H2BK120ac (Active Motif; 39119), H3 (Invitrogen; PA516183), and H3K27ac (Active Motif; 39134). All antibody concentrations are based on the manufacturer guidance. HRP conjugated goat anti-mouse or anti-rabbit secondary antibodies were used for detection with the Pierce ECL western substrate kit (Thermoscientific; 32209). Blots were visualized using a LiCOR odyssey Fc with 10min exposure for chemiluminesence and 30s exposure at 700nm wavelength for the ladder.

### ATAC-seq

Cells were collected at 90% confluence and total cell count was calculated using a Oroflo MoxiZ cell counter. Three tubes of greater than 500,000 for each sample were than crypreserved in our growth media with 10% DMSO added. The samples were than sent to Genewiz for tagmentation (ATAC), library prep and sequencing. These experiments were done in triplicate using three independent cell passages.

### RNA-seq

Cells were grown in a 6-well plate and RNA was collected using 1 mL of Trizol (ThermoFisher; 15596018) according to the manufacturer’s instructions. RNA was isolated using a phenol-chloroform and RNA concentration was measured by nanodrop. The isolated RNA was then flash frozen and sent to Genewiz for library prep and sequencing. These experiments were done in triplicate using three independent cell passages.

### Spike-In ChIP-seq

ChIP-seq was performed in duplicate following the protocol in Chen, H. et al. 2014^33^ with NIH3T3 spike-in chromatin added post sonication. Total chromatin was determined using nanodrop and enough spike-in chromatin was added to constitute 5% of the total chromatin. The same crosslinked and sonicated NIH3T3 chromatin was used as spike-in for all experiments. For immunoprecipitating BRD4 we used the same cell signaling antibody as was used for western blotting, again following the manufacturer guidance for the antibody concentration. These experiments were done in duplicate using two independent cell passages while the same NIH3T3 spike-in chromatin was used for all samples.

### Sequencing Processing Pipeline

All sequencing data had adapters trimmed using Trim galore^43^ followed by being run through fastqc^44^ to check the quality of the initial reads. RNA-seq and ATAC-seq data was aligned using STAR aligner^45^. While the ChIP-seq data was aligned using BWA-MEM2^46^. RNA-seq and ATAC-seq were aligned to the hg19 genome while the ChIP-seq data was aligned to a merged hg19/mm10 hybrid genome with “mm10” added to the mm10 chromosome names (chr1 to mm10chr1). Post alignment, PCR duplicates were marked using Picard tools^47^. Meta plots were created using deeptools2^48^. Genomic overlap fisher exact tests were performed using a custom python script. Sex chromosomal and mitochondrial alignments were removed from the ATAC-seq and ChIP-seq datasets. Peaks for the ChIP-seq and ATAC-seq data were called using Genrich^49^. Fragment pileups were created using FeatureCounts^50^ and used for DESeq2^51^ differential analysis of the ATAC-seq and RNA-seq data in R. Genomic track images were generated using IGV^52^. Quality control information for all sequencing datasets can be found in supplementary table 1.

### Spike-in Normalization using NMF

We developed a novel approach for normalizing ChIP-seq data using spiked-in mouse chromatin to quantify the IP efficiency. Our method assumes that an ChIP experiment contains a mixture of background/noise fragments and real IP fragments, and the fact that spike-in mouse chromatins should in theory have the same underlying IP signals and backgrounds. Therefore, we write the observed fragment counts of the spike-in over n genomic windows of k experiments, including IP as well as inputs, as an n x k matrix Y, and it can be represented by Y = WH, where W is an n x 2 matrix with each column representing the background and the perfect IP signal respectively, and H is a 2 x k matrix representing the proportion of the mixture of IP and background for each of k experiments (up to some scaling factors related to sequencing depth). Using Non-Negative Matrix Factorization (NMF), we can deconvolute Y and estimate W and H. We can then use the estimated H, reflecting the IP efficiency, to normalize the IP experiments on human chromatins. More details of this method can be found in the Supplementary Material. To apply the normalization method, we first created bed files for numerous spike-in (mm10) genomic window sizes (ranging from 500bp to 20kb). We then created count tables for said regions using FeatureCounts then ran a custom R script that uses an NMF model to calculate the correction factor from the spike-in count tables.

### MTT Assay and microscopy

To perform the MTT assay cells were seeded on to a 96-well plate with a matching set seeded on to 10cm plates for cell microscopy. These treatment was staggered between the 96 well plate and the 10cm plate by one hour to ensure that both the MTT assay and microscopy could be done exactly 48 hours post treatment. 5mg/mL PBS MTT (invitrogen; #M6494) in PBS was prepared for the assay. Cells were flushed of media and a 100uL 50/50 mixture of serum free media and MTT solution was added to each well followed by a 3 hour 37C incubation period. After the inculabation 150uL of MTT solvent (4mM HCl/0.1% NP40 in isopropanol) was added to each well. The plate was than wrapped with foil and placed on an orbital shaker for 15 minutes with pipetting of the liquid at 5 and 10 minutes. The plate was analyzed using the SpectraMax iD3 plate reader for 590nm absorbance. Results were than analyzed in prism and a paired t-test was used to compare every condition with the H2BK120R condition During the 3 hour MTT incubation brightfield images of the cells on the 10cm plates were taken using the Keyence BZ-X800.

### Data

The ENCODE^26^ accession and GEO numbers for published datasets used can be found in STAR methods key resources table. Enhancer data for IMR90’s was obtained from the Enhancer Atlas database^24^. All sequencing data generated for this paper has been deposited to GEO at GSE288928.

## Supporting information

Key Resources Table

## Acknowlegements

We thank Christopher A. French for providing the doxycycline inducible BRD4-NUT construct. We also thank David Shechter for giving us access to the Keyence BZ-X800 and his lab for assisting with the microcopy image generation. We also want to thank Jon M. Backer for the MTT assay reagents and protocol. This study was supported by R01 CA155232 to M.J.G., P30 CA013330 (Cancer Center Support Grant) to the Montefiore Einstein Comprehensive Cancer Center, and T32 GM007491 (training grant support) to G.A.H and P.D.R.

## Supplement

**Figure S1.**
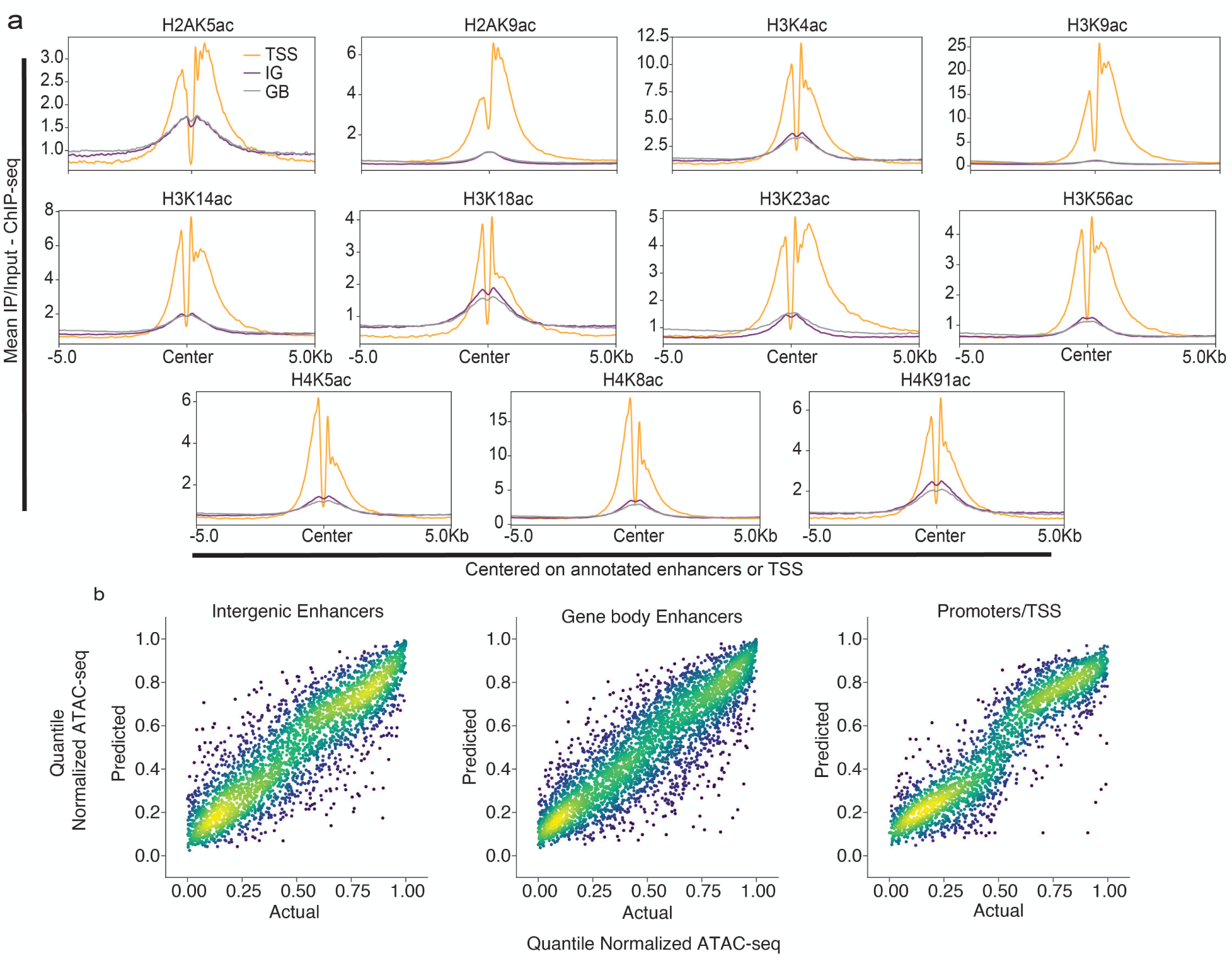
Additional metas and machine learning scatter plots. (a) Additional histone acetylation ChIP-seq meta plots for IGEs(purple), GBEs(grey) and promoters(orange). (b) Scatter plot of actual vs machine learning predicted quantile normalized ATAC-seq signal. IGE(left), GBE(middle) and promters(right). RSQ = .78(IGEs), .76(GBEs) and .83(Promoters), closer to 1 is better. Mean Squared Error = 0.018(IGEs), 0.018(GBEs) and 0.018(Promoters), lower is better.

**Figure S2.**
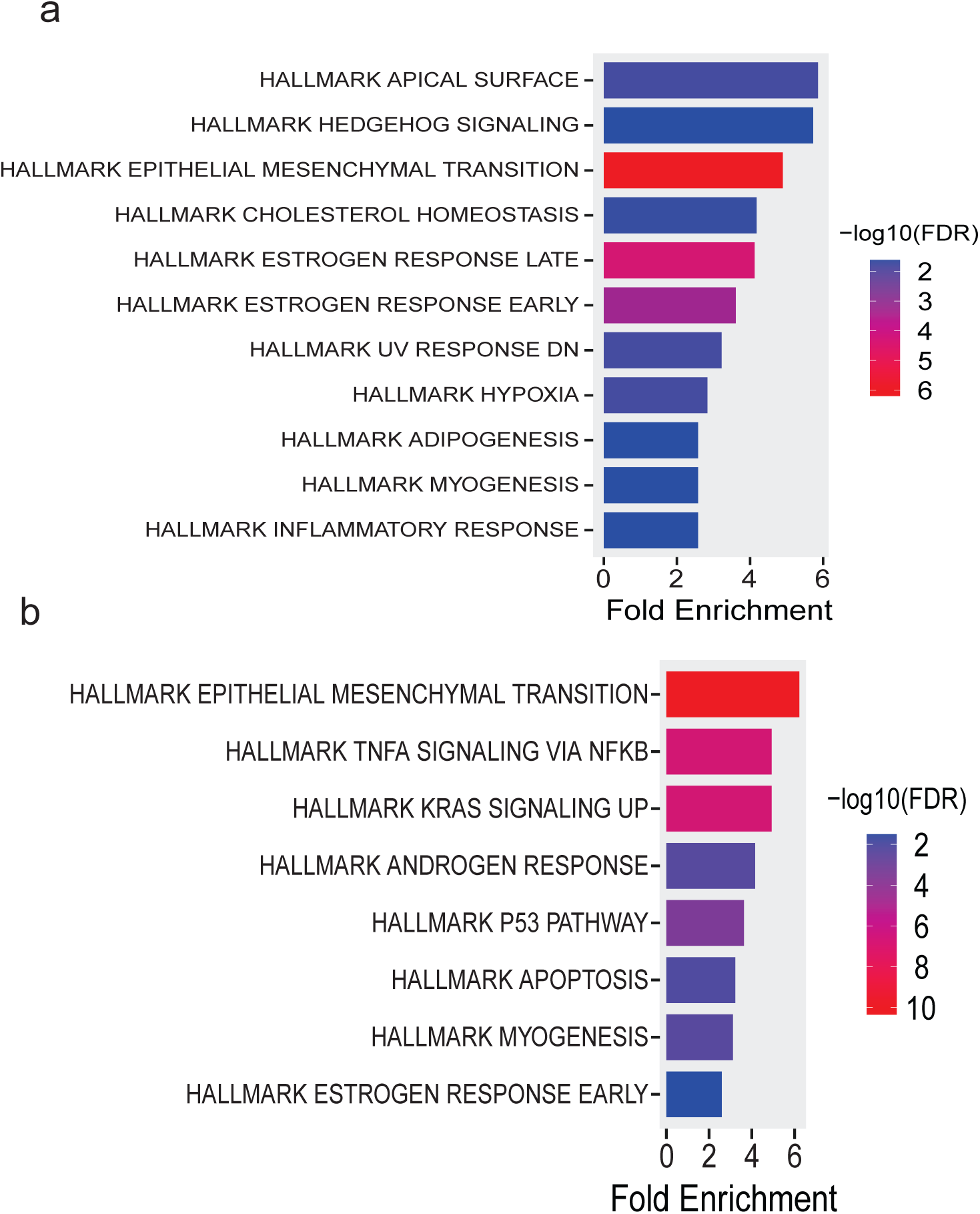
RNA-seq GSEA. GSEA analysis of H2BK120R(a) and H2BK12R(b) RNA-seq DEGs.

**Figure S3.**
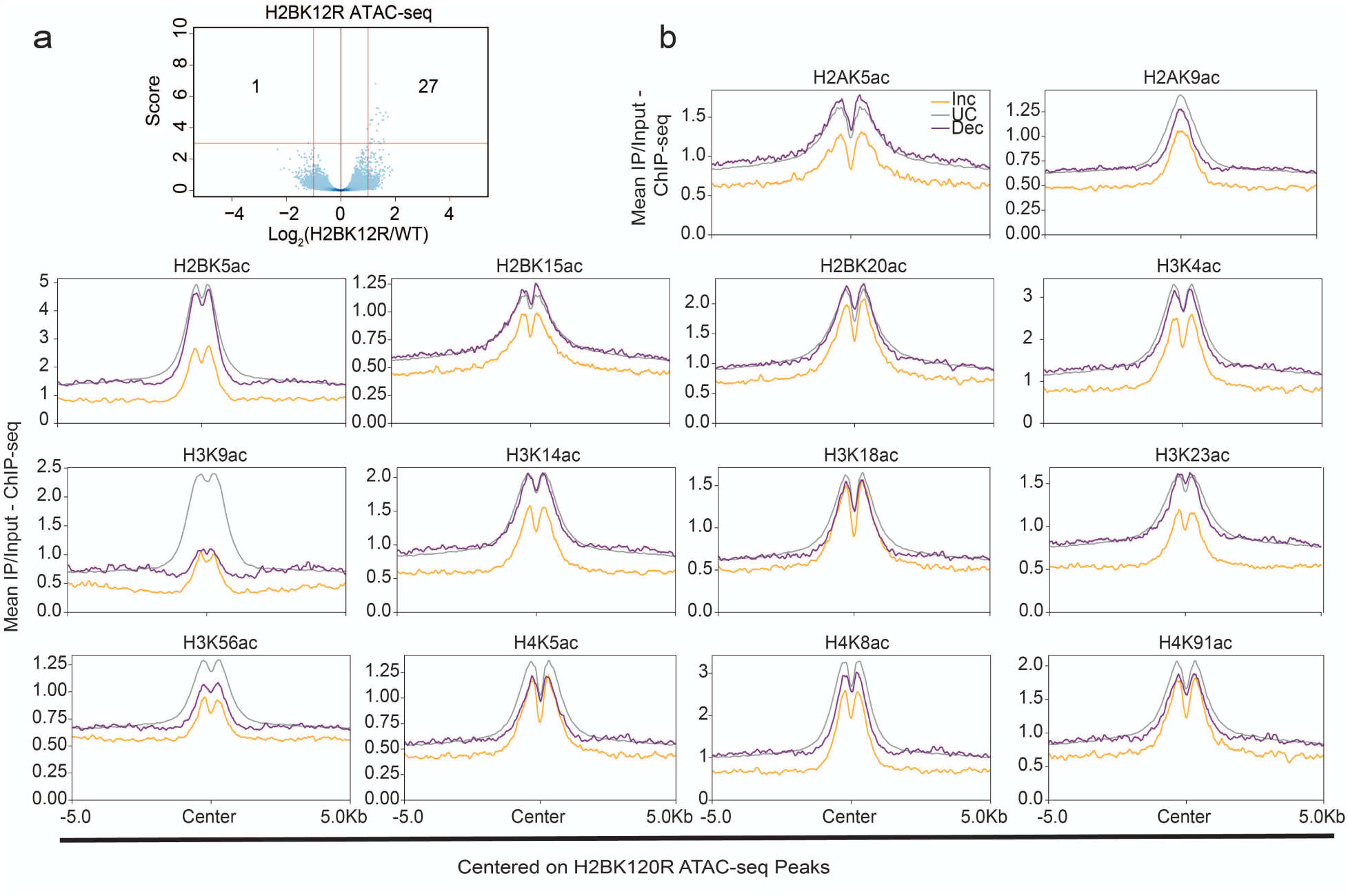
Additional metas and machine learning scatter plots. (a) Volcano plot of H2BK12R ATAC-seq data. (b) Additional histone acetylation ChIP-seq meta plots for H2BK120R ATAC-seq DARs.

**Figure S4.**
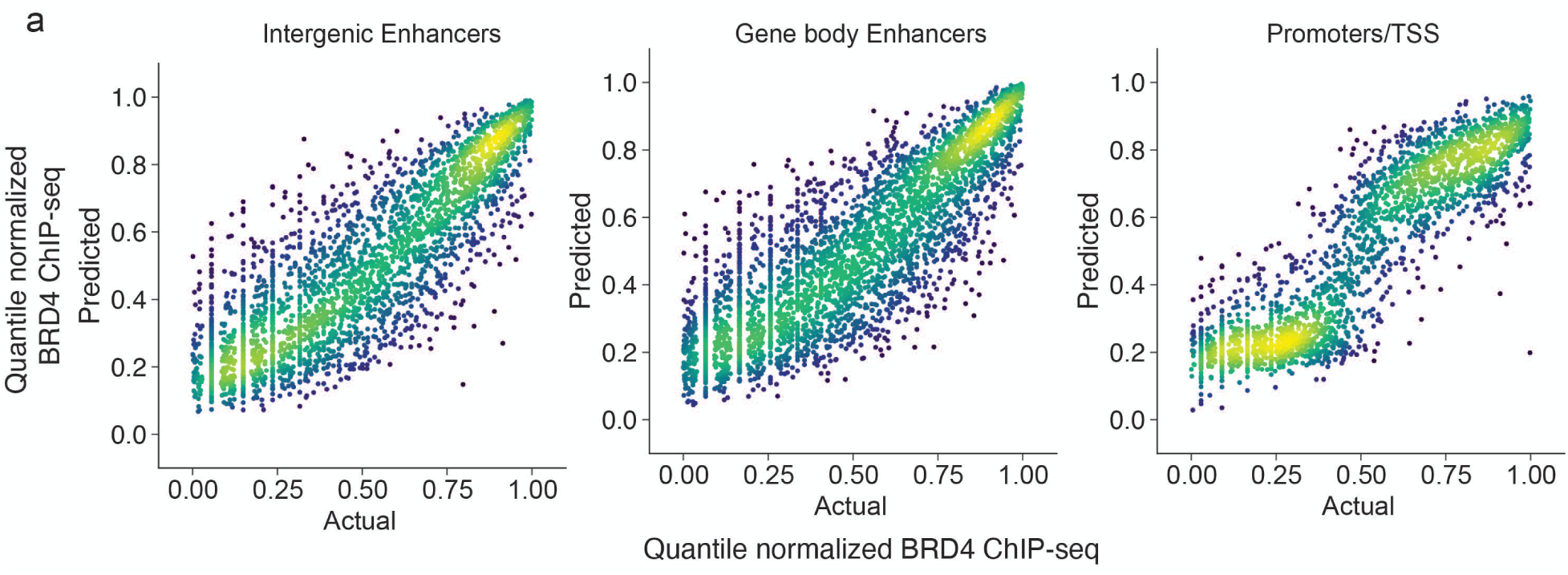
machine learning scatter plots. (a) Scatter plot of actual vs machine learning predicted quantile normalized BRD4 ChIP-seq signal. IGE(left), GBE(middle) and promters(right). RSQ = .76(IGEs), .70(GBEs) and .80(Promoters), closer to 1 is better. Mean Squared Error = 0.020(IGEs), 0.020(GBEs) and 0.016(Promoters), lower is better.

**Figure S5.**
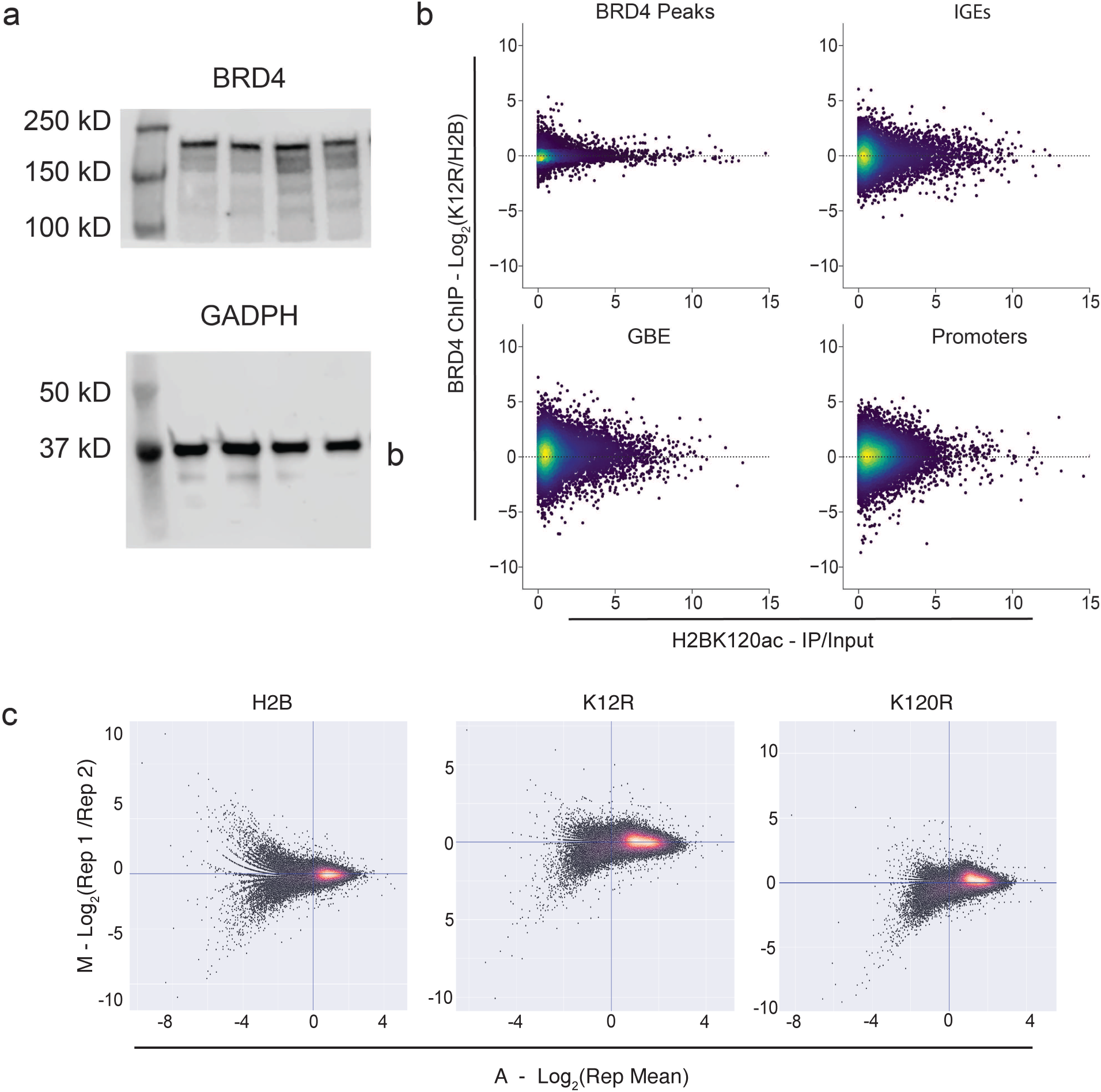
H2BK12ac does not regulate BRD4 Binding. (a) Raw Western blot of Fig 5a (b) Scatter plots of Log2(K12R/H2B)(Left) NMF normalized BRD4 ChIP-seq signal (y-axis) at BRD4 peaks(N=56725, top left), IGEs(N=8484, top right), GBEs(N=9729, bottom left) and promoters(N=15281, bottom right) with IP/Input H2BK120ac ChIP-seq data x-axis. (c) MA Plots comparing NMF normalized BRD4 ChIP-seq the replicates of each condition for 1kb genomic windows. H2B(left), K12R(middle), and K120R (right)

**Figure S6.**
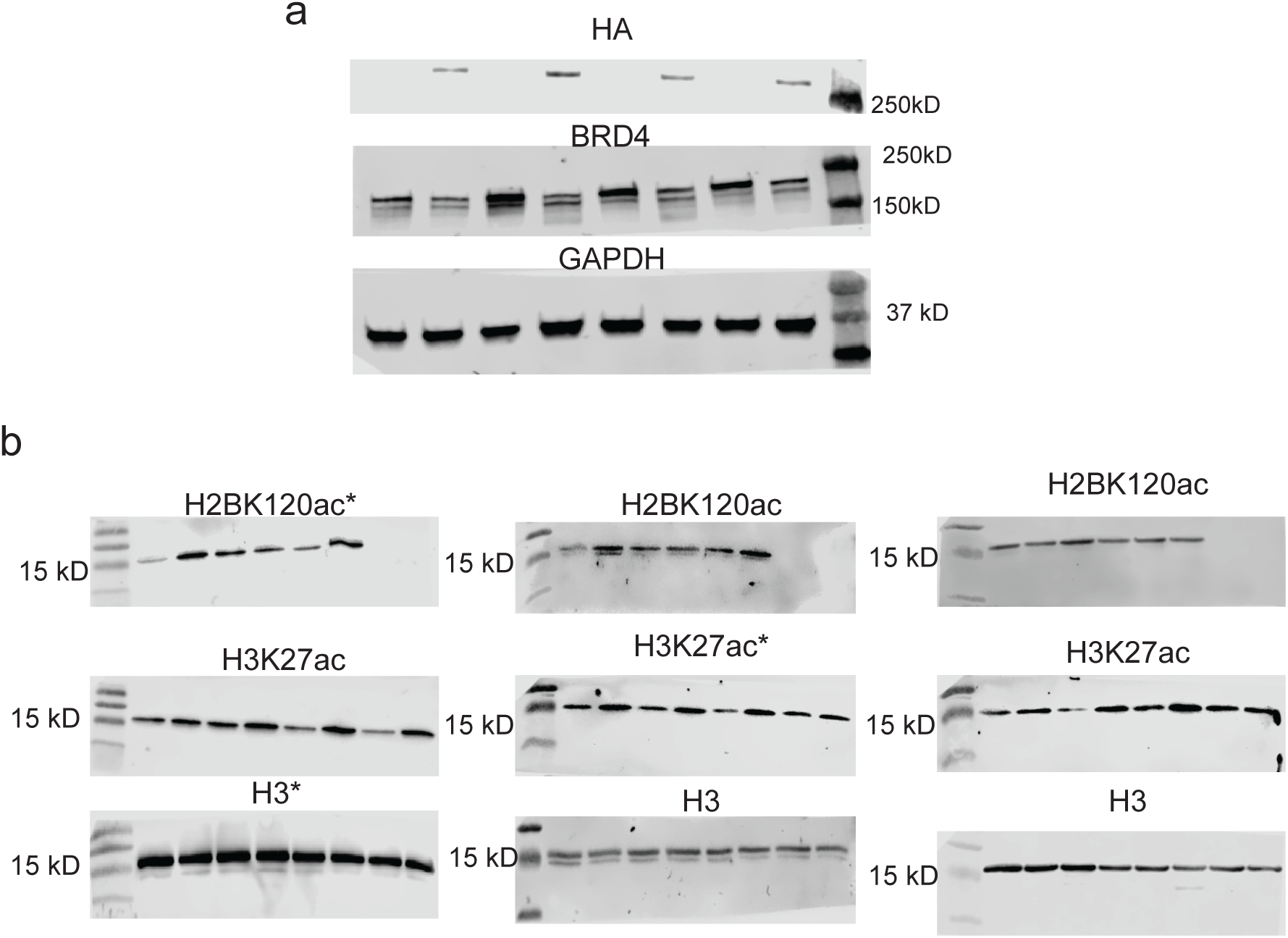
Raw western blots for figure 6 pannels. (a) Raw western blots for Fig 6a left (b) Triplicate raw western blots for Fig 6a right. * Indicates blot used in main figure.

